# Promoter-Driven Modulation of Flagellin Expression and Motility in *Escherichia coli*

**DOI:** 10.1101/2025.01.31.635866

**Authors:** Karima Bharati Bayana, Xinyue Wang

## Abstract

Synthetic biology offers powerful tools to engineer biological systems for diverse applications. However, key challenges persists before achieving real-world applications like environmental bioremediation or therapeutic microrobots for targeted drug delivery. This study aimed to precisely control bacterial movement by modulating gene expression using engineered promoters in Escherichia coli. We focused on Escherichia coli, a model organism, and manipulated its motility by engineering the expression of flagellin, a crucial protein for bacterial movement. To achieve this, specific genetic promoters were employed to regulate the production of flagellin, thereby dictating the movement capabilities of these bacteria. The promoters enabled targeted adjustments to flagellin expression, which in turn allowed for the enhancement or suppression of bacterial locomotion. Interestingly, the relationship between promoter design parameters and gene expression levels was non-linear, highlighting complex underlying dynamics. Optimal bacterial motility occurred at 30°C, illustrating the influence of environmental factors. Our findings demonstrate the ability to effectively regulate complex microbial phenotypes like motility using genetic engineering strategies. The results not only extend our understanding of bacterial gene regulation but also highlight the transformative potential of synthetic biology in creating functional and adaptable microbial phenotypes for diverse biotechnological applications.

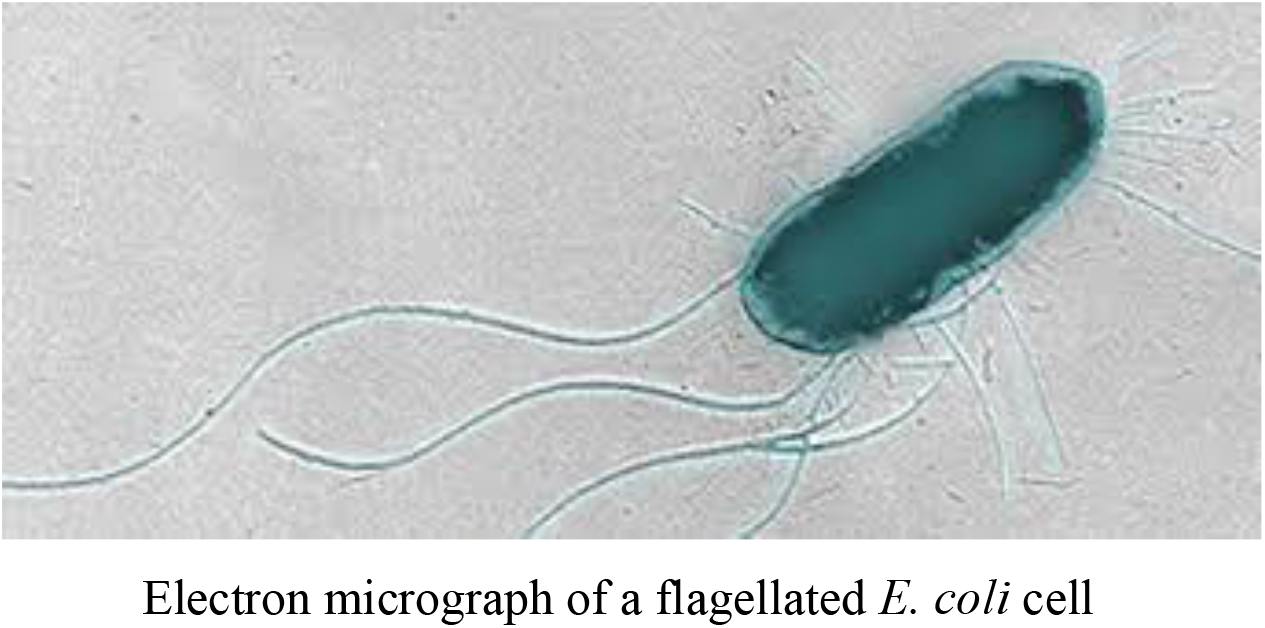

## Introduction

In the rapidly evolving field of synthetic biology, the manipulation of bacterial motility offers innovative solutions to some of the most challenging problems in various sectors. At the core of this manipulation is the ability of bacteria such as Escherichia coli to move using a complex motor system powered by flagella. These structures act as tiny propellers, driven by rotary engines made from protein complexes, enabling bacteria to navigate through various environments (Guo and Liu, 2022). This natural phenomenon has significant potential for applications ranging from biodesalination, which could help address global freshwater shortages, to the development of bacterial microrobots that could deliver drugs directly to diseased cells (López-Igual et al., 2019).

The flagellar system in E. coli is powered by a molecular motor that requires the coordinated expression of more than 50 genes, including fliC, which codes for the structural component of the flagellum known as flagellin (Kakkanat et al., 2015). Regulation of flagellar synthesis is tightly controlled by a hierarchical cascade of genetic elements, which presents an ideal model for synthetic regulation using engineered promoters.

The control of this motility is heavily influenced by promoters, which are regions of DNA that is responsible for the construction and operation of flagella. Promoters act as critical components for controlling gene expression. Synthetic biologists have learned to modify these promoters to adjust the speed, efficiency, and responsiveness of bacterial movement to external stimuli. This study employs two specific variants from the Anderson promoter library, which includes a range of constitutive promoters with variable expression strengths, facilitates the modulation of gene expression without the reliance on additional regulatory proteins, thus offering a streamlined approach to gene regulation (Rytter et al., 2014). The promoters J23101 and J23106, are used to examine the correlation between promoter strength and gene expression. In addition, we also incorporates the lacI/IPTG-inducible promoter, BBa_R0011, to achieve controllable expression of target genes in response to exogenously supplied IPTG, allowing for the detailed examination of gene expression dynamics in response to environmental cues(Marbach and Bettenbrock, 2012). This dual promoter approach is used to regulate the expression of the gfp gene, facilitating the real-time monitoring of promoter activity, and the fliC gene, which directly impacts the motility of Escherichia coli.

The aim of our study revolves around utilizing these synthetic promoters to regulate the expression of two key genes: GFP and fliC. The GFP serves as a reporter gene, allowing for the visualization and quantification of gene expression, whereas the fliC gene directly influences bacterial motility. By integrating these genetic constructs into E. coli strains, we can explore the efficacy of different promoters, thereby gaining valuable insights into the dynamic regulatory networks of bacterial gene expression.

Furthermore, the broader implications of this study extend to potential applications such as the development of bacterial microrobots for targeted drug delivery and biodesalination processes using genetically engineered bacteria to remove salt from seawater, thus addressing global freshwater shortages (Johnson and Li, 2018; Smith et al., 2020).

## Methods

All methods were carried out as stated in the lab manual (Colloms, 2024), with the only exception being a change in the time interval for measuring OD values.

### Results Plasmid Design

To investigate the use of synthetic promoters for controlling gene expression in Escherichia coli, focusing on the GFP reporter gene and fliC, which affects bacterial motility. Plasmids were constructed with GFP and fliC genes under the control of the J23101 and J23106 promoters inserted between the EcoRI and the XbaI sites from the Anderson promoter library and the inducible promoter BBa_R0011. As part of our verification process, we performed restriction digests on our plasmid samples using HindIII, alongside other enzymes like EcoRI and PstI, to map out the restriction sites accurately. The digests were resolved on an agarose gel to visualize the DNA fragments, the absence of HindIII restriction sites in our plasmid constructs was confirmed as shown in the agarose gel electrophoresis image (Figure 3).

**Figure 1.**
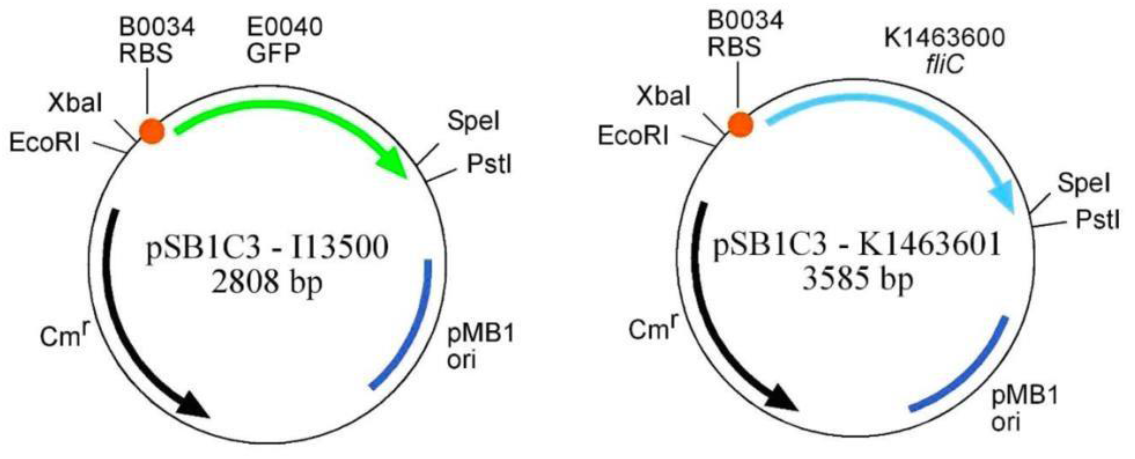
Plasmid Maps without promotors for GFP and FliC expression.

**Figure 2:**
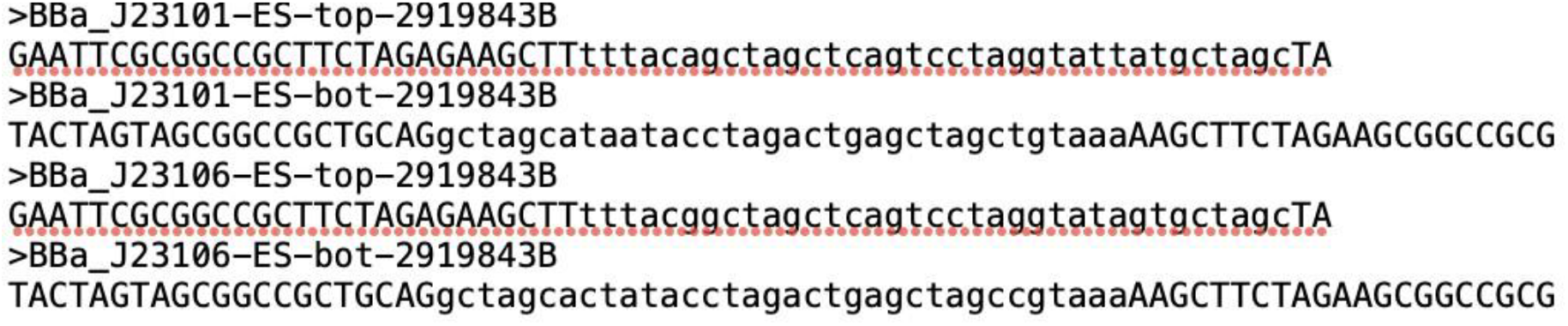
Sequences of Synthetic Promoters BBa_J23101 and BBa_J23106. This figure displays the nucleotide sequences of synthetic promoters BBa_J23101 and BBa_J23106, highlighting their forward and reverse orientations. The sequences illustrate transcription start sites and core promoter elements, emphasizing regions critical for promoter activity.

**Figure 3:**
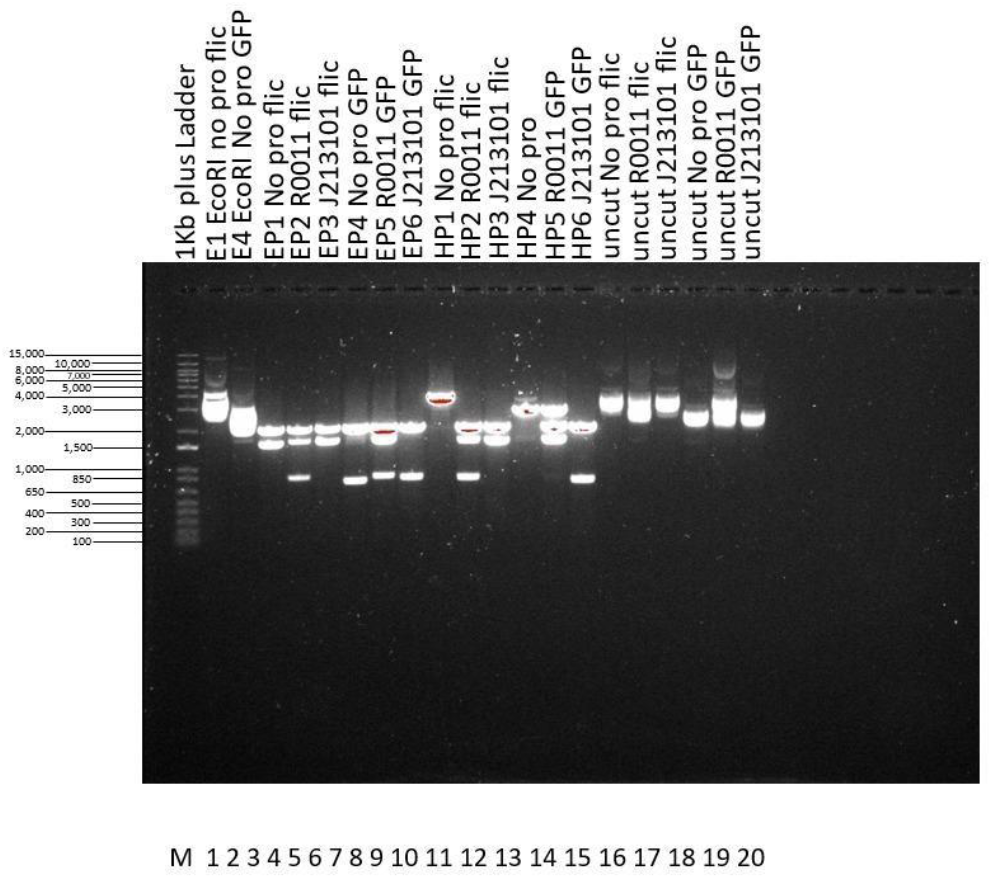
Restriction Digests of Plasmids Containing GFP and fliC Constructs. This gel image displays an ethidium bromide-stained agarose gel, showcasing restriction digests of GFP and fliC plasmids. Lane M contains a 1Kb plus DNA ladder. Lanes 1-15 illustrate digests with EcoRI, PstI, and HindIII across different promoter variants, while lanes 16-20 display uncut plasmid controls, indicating supercoiled DNA. Each lane is labeled according to the enzyme and plasmid type.

### Quantitative analysis of GFP production from Plasmids

In this experiment, we conducted quantitative analyses to evaluate the promoter strengths of J23101, J23106, and R0011 in response to varying concentrations of isopropyl β-D-1thiogalactopyranoside (IPTG) and to assess the inducibility of the R0011 promoter within the Escherichia coli strain DS941 Z1 through GFP fluorescence measurements.

Firstly, to evaluate the inducibility of the GFP gene under the R0011 promoter in the DS941 Z1 strain by IPTG, we conducted fluorescence measurements. We grew DS941 Z1 cultures with a GFP-bearing plasmid under the R0011 promoter, induced one group with IPTG, and left the other uninduced. We measured fluorescence every 30 minutes over four hours to track GFP production.

The IPTG-induced culture showed a marked increase in fluorescence, demonstrating the R0011 promoter’s strong inducibility and tight regulation of GFP expression in this strain (figure 4).

**Figure 4:**
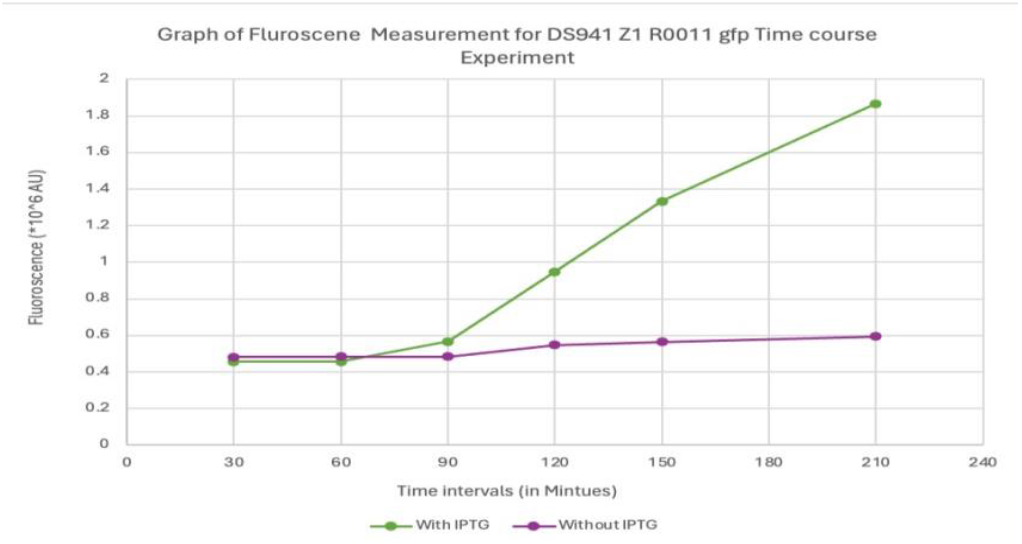
Time-Course Fluorescence Measurement for DS941 Z1 R0011 GFP Expression. This graph illustrates the fluorescence intensity of GFP in DS941 Z1 cells under the R0011 promoter, measured over a 4-hour period. The green line represents cells induced with IPTG, showing a significant increase in fluorescence over time. The purple line depicts uninduced cells, maintaining low fluorescence throughout the experiment.

The uninduced control maintained stable fluorescence levels, indicating minimal background expression from the R0011 promoter.

These findings confirm the R0011 promoter’s capability to precisely control GFP expression in response to IPTG, making it suitable for applications requiring exact gene expression control.

We also assessed the impact of IPTG on the growth rate of Escherichia coli harboring GFPexpressing plasmids by monitoring optical density at 600 nm (OD600) over time. We cultured E. coli in LB medium, dividing it into two groups: one that we induced with IPTG and another left uninduced. We recorded OD600 readings every 30 minutes for a total of 4 hours to track cell growth. Both groups displayed similar growth patterns, robustly increasing over the experiment’s duration. An initial lag phase lasting about 60 minutes preceded a log phase characterized by rapid growth (figure 5).

**Figure 5:**
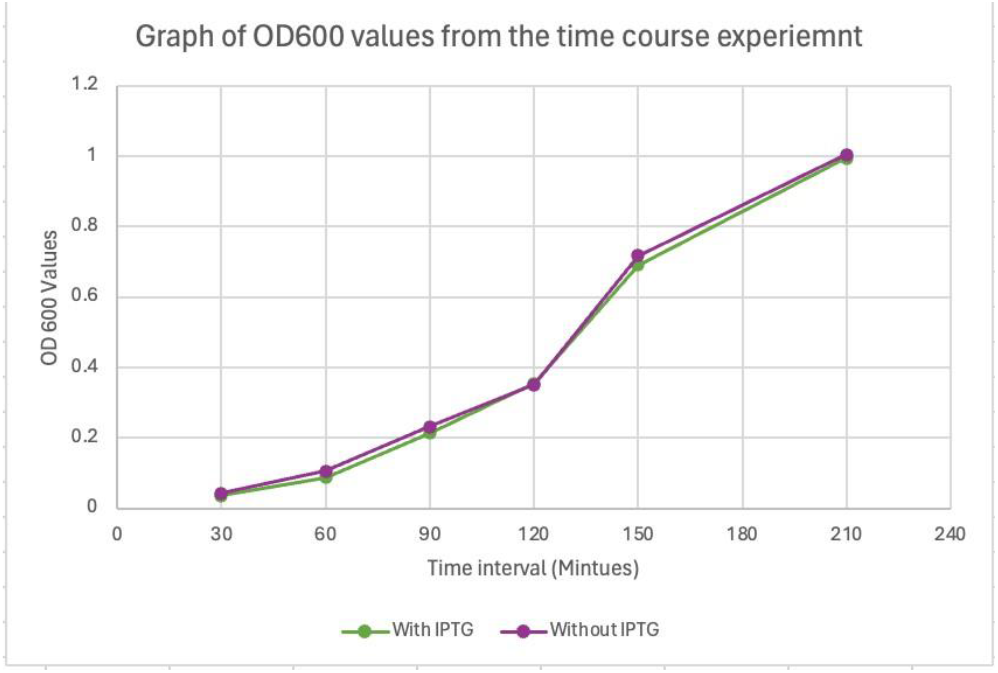
Growth Kinetics of E. coli with and without IPTG Induction. This graph presents OD600 measurements taken every 30 minutes over a four-hour period, tracking the growth of E. coli carrying GFPexpressing plasmids. The green line represents cells induced with IPTG, while the purple line shows uninduced cells, illustrating similar growth patterns for both conditions.

The presence of IPTG did not significantly affect the overall growth rate, as the overlapping curves indicated. This observation suggests that IPTG induction of GFP production does not impose a significant metabolic burden that could disrupt growth dynamics under these testing conditions. Our data confirm that IPTG induction does not adversely impact the growth of E. coli cultures expressing GFP, ensuring that the expression system remains viable for further experiments without compromising bacterial health or proliferation.

Furthermore, to evaluate the relationship between cell concentration and GFP expression, we measured the fluorescence intensity of GFP across a range of cell dilutions. We recorded the fluorescence in arbitrary units (AU) to assess the level of GFP expression relative to the cell density (Figure 6). Both curves consistently showed that GFP fluorescence intensity decreased with increasing dilution, indicating a proportional relationship between cell density and GFP expression. This proportionality suggests that GFP production per cell remains constant across different cell densities.

**Figure 6:**
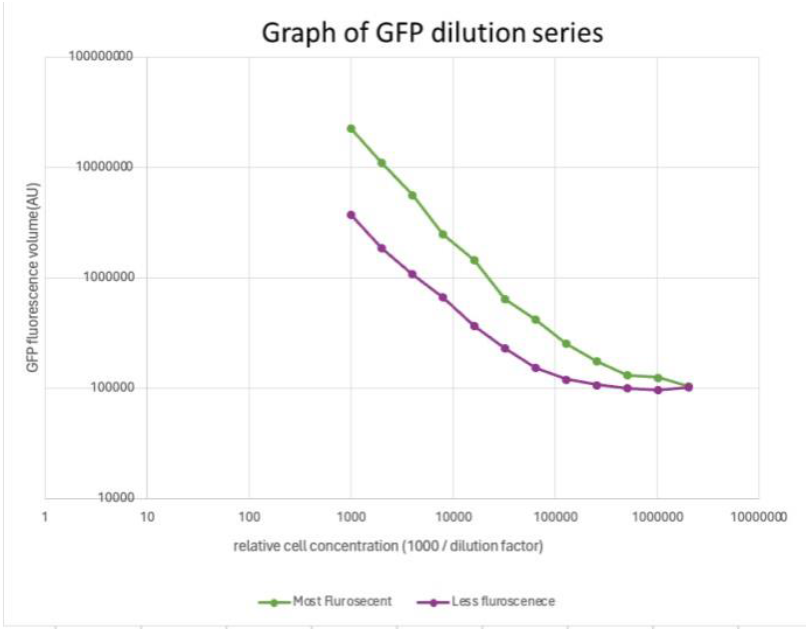
GFP Fluorescence Intensity Across Dilution Series. This graph shows GFP fluorescence measured in arbitrary units (AU) against relative cell concentration (1000/dilution factor) on a logarithmic scale. The green line represents samples with higher initial GFP expression, while the purple line represents those with lower expression, both depicting a decrease in fluorescence with increasing dilution.

The distinct separation between the two curves throughout the dilution series highlighted the impact of initial expression levels on the observable fluorescence, emphasizing variability in promoter strength or plasmid copy number among different samples. The data clearly demonstrated that GFP fluorescence correlates with cell concentration, validating the use of GFP as a quantitative marker in cellular assays. The variations in expression levels captured in the graph also underscore the importance of standardizing cell density in experiments using fluorescent reporters to ensure accurate interpretation of expression data.

To compare the strength and inducibility of various promoters (J23101, J23106, and R0011) in GFP expression systems, we measured fluorescence across different IPTG concentrations. We grew cultures with plasmids containing different promoter constructs and induced them with IPTG ranging from 0 mM to 1 mM. We quantified GFP production through fluorescence measurements, offering a direct assessment of promoter activity under each tested condition (figure 7).

**Figure 7:**
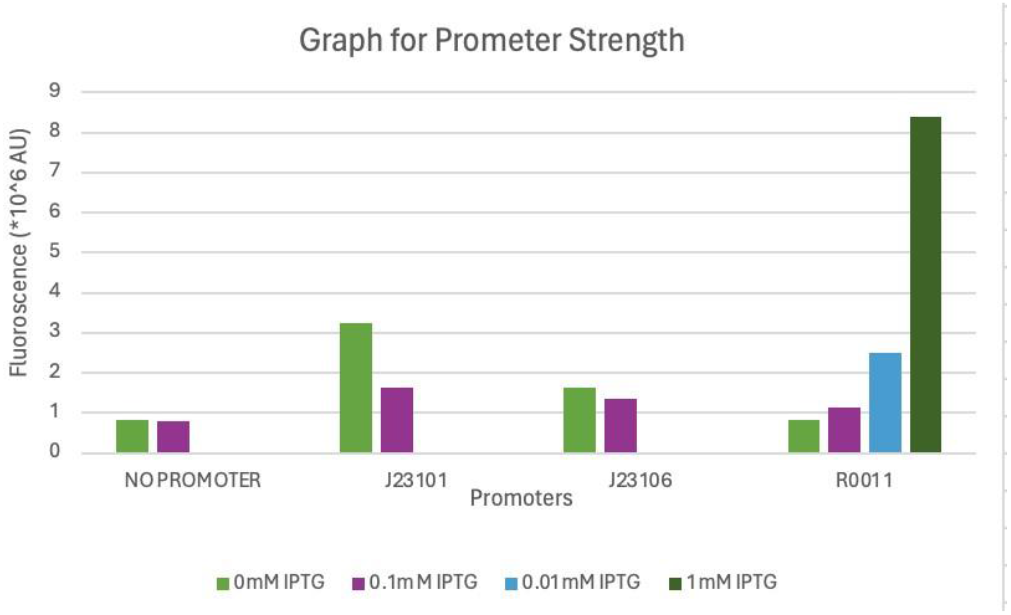
Fluorescence response of GFP expression driven by different promoters (No Promoter, J23101, J23106, R0011) at various IPTG concentrations (0 mM, 0.01 mM, 0.1 mM, 1 mM). Fluorescence intensities are shown in arbitrary units (AU × 10^6) to compare promoter strength and inducibility.

The control with no promoter displayed minimal fluorescence at all IPTG concentrations, setting a baseline for background fluorescence. The J23101 promoter showed moderate fluorescence, with slight responsiveness to IPTG, indicating its low inducibility. The J23106 promoter demonstrated similar behavior to J23101 but with slightly lower fluorescence intensities, suggesting a weaker promoter under these experimental conditions. Conversely, the R0011 promoter exhibited a dramatic increase in fluorescence at higher IPTG concentrations (1 mM), confirming its strong responsiveness to IPTG and high inducibility.

This data highlights significant variability in promoter activity, with R0011 emerging as the most inducible by IPTG, making it ideal for applications requiring tight regulation of gene expression. In contrast, J23101 and J23106, with their lower and more consistent activities, are better suited for applications needing steady expression without the necessity for induction control.

### Restoring and Controlling Motility in E. coli

The motility assays were performed to evaluate the ability of these promoters to restore and control motility in the fliC-deficient strains. In evaluating the motility of E. coli strain DS941, we found significant variation depending on the promoter used and the presence of IPTG as seen in figure 8. The promoter J23101 with 0.1 mM IPTG greatly enhanced motility, achieving a spread of approximately 6 cm, which was near the level of the motile control at 7 cm. Without IPTG, J23101’s effectiveness dropped to about 3 cm. In contrast, promoter J23106 only allowed for a motility of 1 cm, indicating a lower efficiency under our test conditions.

**Figure 8:**
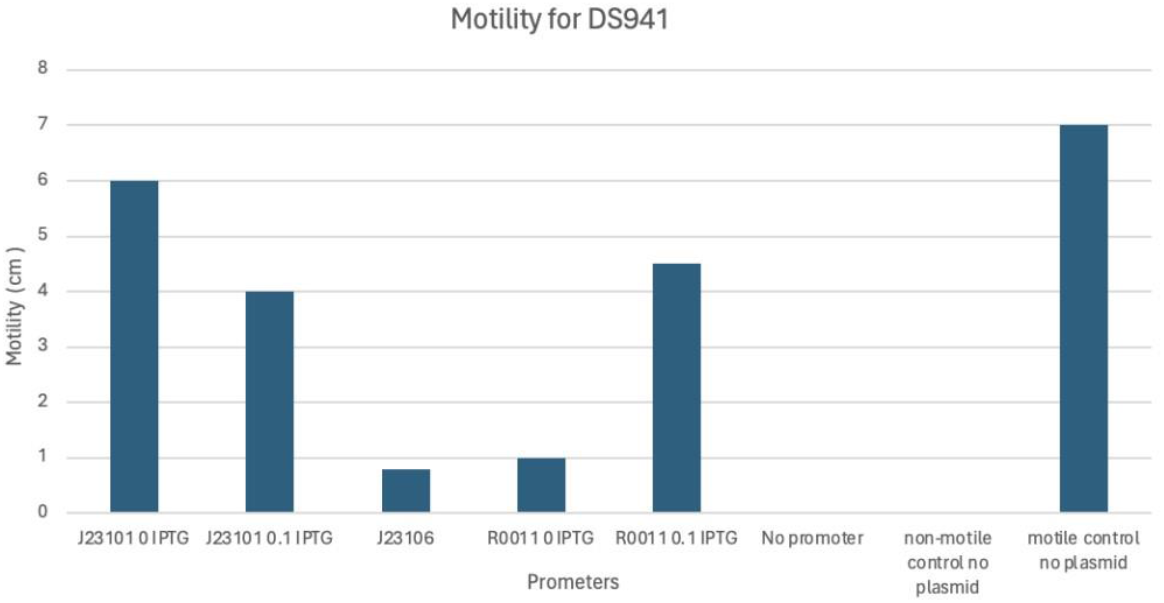
Bar graph representing motility (in cm) of strain DS941 under various conditions. The x-axis shows different promoters (J23101 and R0011) with or without IPTG induction, along with controls lacking a promoter, and motility controls with and without a plasmid.

The IPTG-inducible promoter R0011 demonstrated conditional behavior; it supported a motility of 4 cm with IPTG but showed negligible movement without it, akin to the non-motile control. This highlights the importance of IPTG in activating the R0011 promoter for motility in E. coli.

In the evaluation of promoter strength based on fluorescence output, the recorded data illustrates a nonlinear relationship between the numerical promoter strength index and fluorescence intensity, measured in arbitrary units (AU) (figure 9). At a promoter strength index of 0.5, the fluorescence output was approximately 150,000 AU. When the promoter strength was increased to 1.0, the fluorescence unexpectedly decreased to around 125,000 AU. However, further increase in promoter strength to 1.5 resulted in a moderate increase in fluorescence to about 175,000 AU. Notably, the highest fluorescence observed was approximately 325,000 AU at a promoter strength of 2.5, indicating a significant upsurge in promoter activity compared to lower strength indices. This pattern suggests that promoter efficacy may be influenced by factors beyond mere numerical increases in promoter strength.

**Figure 9:**
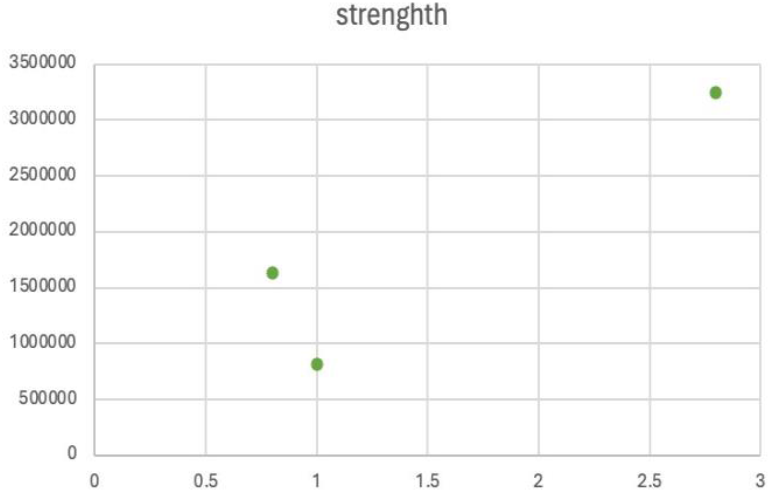
Fluorescence Intensity by Promoter Strength. This scatter plot displays the relationship between promoter strength (ranging from 0 to 3) and fluorescence intensity in arbitrary units (AU) for E. coli. Data points at promoter strengths of 0.5, 1.0, 1.5, and 2.5 highlight the non-linear response of fluorescence output to increases in promoter strength.

Furthermore, we assessed the motility of the E. coli strain DS941 and its plasmid-free mutant, DS941ΔfliC, across various temperatures 37°C, 30°C, and 25°C. We observed significant motility in the DS941 strain at all temperatures, noting a decline in motility with decreasing temperature: 8.5 cm at 37°C, 5.4 cm at 30°C, and 3.2 cm at 25°C (figure 10). Conversely, the DS941ΔfliC mutant exhibited no motility at any of the temperatures, confirming the importance of the plasmid in motility expression.

**Figure 10:**
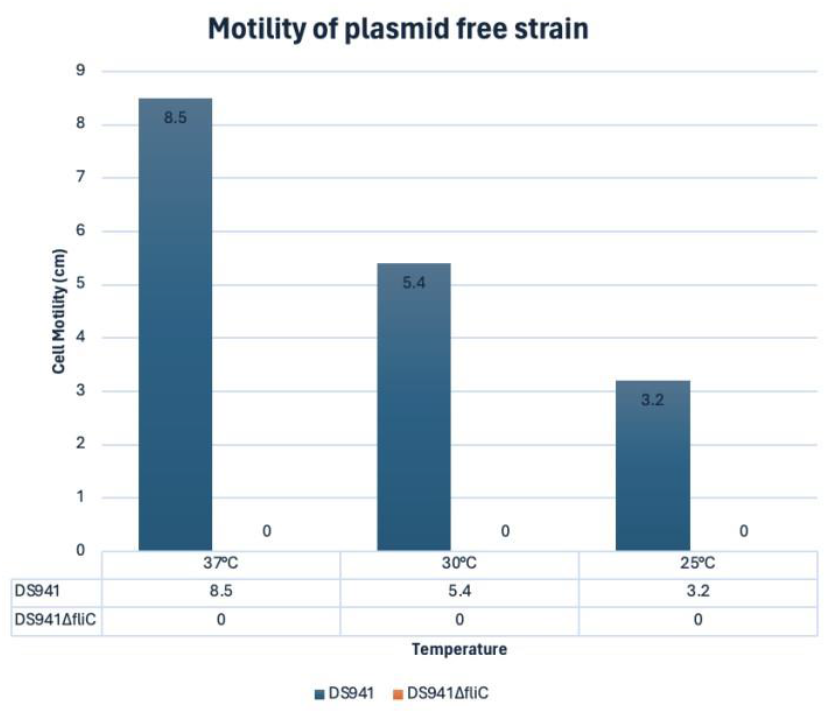
Motility Comparison of E. coli Strains DS941 and DS941ΔfliC at Various Temperatures. This bar graph illustrates the motility, of the E. coli strain DS941 and its plasmid-free mutant DS941ΔfliC across temperatures of 37°C, 30°C, and 25°C.

### Overview of Fluorescence and Motility Assays

The experiments conducted across various groups in the class explored the effects of different IPTG concentrations on the fluorescence volumes in E. coli strains, both Z1 and non-Z1 backgrounds, with different promoters inserted upstream of the GFP gene. A box whisker plot was used to summarize these fluorescence measurements across IPTG concentrations of 0.0 mM, 0.01 mM, 0.1 mM, and 1.0 mM (Figure 11).

**Figure 11:**
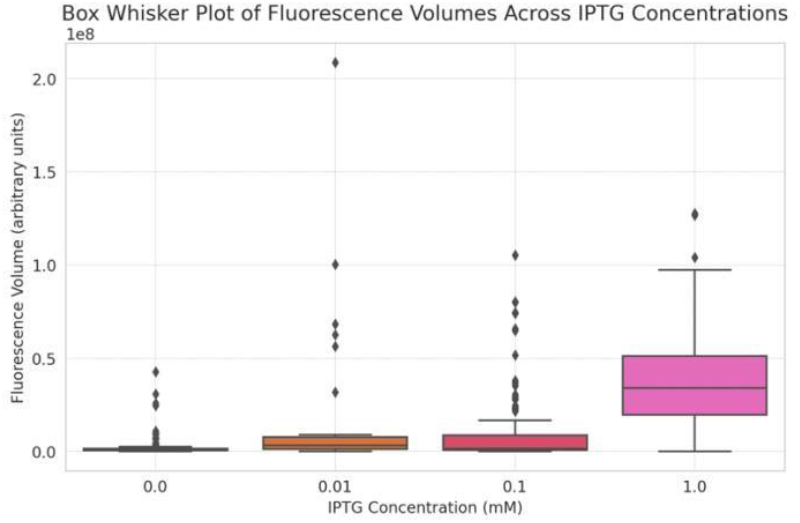
Box-and-whisker plot illustrating the distribution of fluorescence volumes in response to various IPTG concentrations (0.0 mM, 0.01 mM, 0.1 mM, and 1.0 mM). The central line in each box represents the median, the boxes cover the interquartile range (IQR), and the whiskers extend to the farthest points that are not considered outliers. Data points outside of the whiskers are plotted individually as diamonds, indicating outliers.

Our experiment revealed clear variations in E. coli motility at different temperatures, based on data averaged across the entire class which can been seen in figure 12. At 25°C, the average motility diameter was 1.5 cm. The motility significantly increased at 30°C, where we observed the highest average diameter of 2.3 cm. However, at 37°C, the motility decreased to an average diameter of 1.2 cm. These results suggest that E. coli exhibits optimal motility at 30°C under the conditions tested in our class experiments.

**Figure 12:**
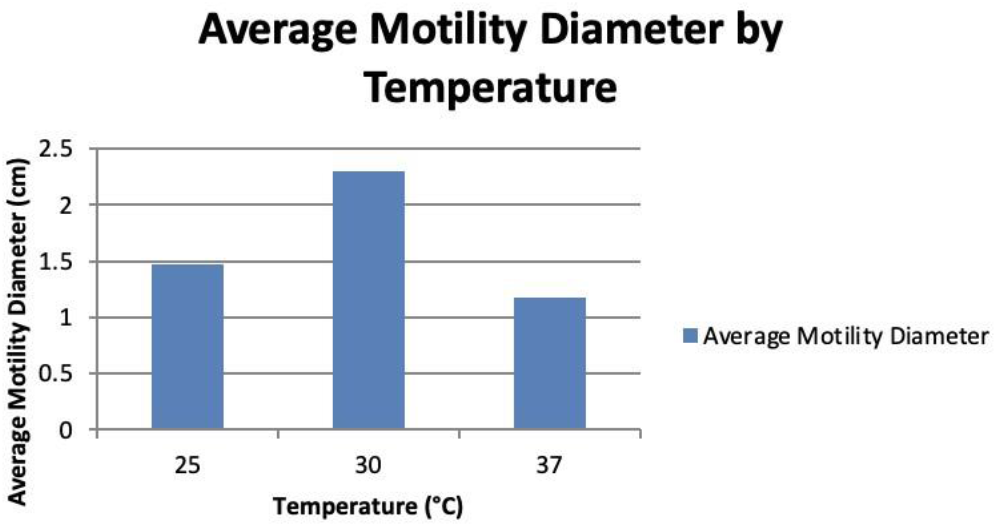
Average Motility Diameter by Temperature. The bar chart shows the mean motility diameter of E. coli at 25°C, 30°C, and 37°C. The highest motility occurs at 30°C, suggesting this temperature is optimal for bacterial movement.

The findings from this study explain the successful application of synthetic promoters in the regulation of gene expression and modulation of bacterial motility in the model organism Escherichia coli. The utilization of constitutive promoters J23101 and J23106, facilitated the attainment of variable levels of green fluorescent protein (GFP) expression and restoration of motility in the fliC-deficient strain. Notably, the J23101 promoter exhibited a potent capacity to recover motility, achieving a radial spread comparable to the motile control strain. Conversely, the implementation of the inducible promoter R0011, in conjunction with the inducer isopropyl β-D-1-thiogalactopyranoside (IPTG), demonstrated precise regulation over gene expression and motility dynamics, with the presence of IPTG eliciting substantial augmentation in these parameters.

## Discussion

The aim of this study was to investigate the use of synthetic promoters for modulating gene expression and motility in the model organism Escherichia coli. The results of this study provide insightful contributions to the understanding of synthetic promoter control over Escherichia coli motility and gene expression, reflecting both the anticipated and novel aspects of bacterial behavior under engineered genetic regulation. The constitutive J23101 promoter from the Anderson library facilitated substantial restoration of motility in the fliC-deficient strain, nearly matching the wildtype control. This aligns with previous characterizations of J23101 as a strong constitutive promoter suitable for robust gene expression (Mutalik et al., 2013). In contrast, the relatively weaker J23106 promoter was less effective at restoring motility, consistent with its lower transcriptional output reported in the literature (Rytter et al., 2014).

The IPTG-inducible R0011 promoter exhibited exquisite control over gene expression, with IPTG induction leading to marked increases in GFP fluorescence and motility. This tight regulation is expected based on the well-established lacI/IPTG system for inducible gene expression in E. coli (Lutz, 1997). The notable increase in GFP expression upon IPTG induction in our study confirms the promoter’s effective inducibility, which is consistent with the findings of Guo and Liu (2022), who noted similar responsiveness in their systems. Importantly, the lack of IPTG resulted in negligible expression, confirming the stringent repression of this promoter in its uninduced state.

Conversely, the variable expression observed with the J23101 and J23106 promoters deviates slightly from Rytter et al. (2014), who suggested more consistent outcomes with the Anderson library promoters. Our results suggest that cellular context, such as promoter placement and chromosomal environment, could significantly affect the behavior of synthetic promoters, indicating a complex interaction between promoter architecture and the cellular transcription machinery.

The temperature dependence of motility, with optimal movement at 30°C, matches expectations from prior work examining the effects of temperature on flagellar assembly, motor function, and chemotaxis in E. coli (Darnton et al., 2007). Thermal stress at higher temperatures like 37°C likely disrupts proper flagellar function.

This study’s findings were mirrored across different experimental setups involving classmates working with other promoter variants and in different E. coli strains, such as the Z1 and nonZ1 backgrounds. Notably, variations in promoter performance between these backgrounds underscored the impact of genetic context on promoter efficacy, which is a critical consideration for synthetic biology applications.

The non-linear relationship between promoter strength and gene expression highlights an area for further research. Investigating the mechanisms behind this phenomenon could involve the use of computational modeling to predict promoter behavior or the engineering of novel promoter architectures to enhance predictability and efficiency. Additionally, expanding the range of environmental conditions tested could provide deeper insights into the robustness of these synthetic systems under diverse operational conditions.

Collectively, our findings demonstrate the power of synthetic biology tools in precisely controlling complex phenotypes like bacterial motility. However, key challenges remain before realizing applications like bacterial microrobots for drug delivery. These include developing robust navigation systems, integrating synthetic sensing/response circuits, and optimizing for complex in vivo environments. For biodesalination, motility control must be coupled with expression of desalinating proteins/transporters tuned to specific salt gradients (Xu et al., 2021).

This study not only enhances our understanding of synthetic promoter systems in controlling bacterial motility and gene expression but also sets the stage for the use of synthetic biology in developing complex, responsive, and safe biological systems for environmental and therapeutic applications. Future experiments should focus on refining promoter design, exploring the interaction of genetic circuits with host physiology, and testing these engineered systems in real-world scenarios to fully harness their potential.

